# Suppression of non-homologous end joining does not rescue DNA repair defects in Fanconi anemia patient cells

**DOI:** 10.1101/151472

**Authors:** Supawat Thongthip, Brooke A. Conti, Francis P. Lach, Agata Smogorzewska

## Abstract

Severe cellular sensitivity and aberrant chromosomal rearrangements in response to DNA interstrand crosslink (ICL) inducing agents are hallmarks of Fanconi anemia (FA) deficient cells. These phenotypes have previously been ascribed to inappropriate activity of non-homologous end joining (NHEJ) rather than a direct consequence of DNA ICL repair defects. Here we used chemical inhibitors, RNAi, and Clusterd Regularly Interspaced Short Palindromic Repeat (CRISPR)-Cas9 to inactivate various components of NHEJ in cells from FA patients. We show that suppression of DNA-PKcs, DNA Ligase IV and 53BP1 is not capable of rescuing ICL-induced proliferation defects and only 53BP1 knockout partially suppresses the chromosomal abnormalities of FA patient cells.

## INTRODUCTION

Fanconi anemia (FA) is a rare, genetically heterogeneous disease that clinically presents as an array of congenital abnormalities, bone marrow failure and predisposition to cancers (Auerbach, 2009). Thus far, 21 genes have been identified as mutated in FA patients (FANCA-FANCV) (Mamrak et al., 2016). The defining characteristics of FA patient cells are cellular hypersensitivity and chromosomal instability upon exposure to DNA interstrand crosslink (ICL)-inducing agents such as mitomycin C (MMC) and diepoxybutane (DEB) (Auerbach and Wolman, 1976). FANC proteins are necessary for the repair of ICLs (reviewed in Kottemann and Smogorzewska, 2013) and also function in protecting replication forks under stress (Schlacher et al., 2011; Schlacher et al., 2012)

Recent studies suggest that the defects in cells deficient in the FA pathway stem from the engagement of the non-homologous end joining pathway (Adamo et al., 2010; Pace et al., 2010). It was proposed that, in the absence of proper repair of ICLs, the NHEJ pathway aberrantly repairs DNA double-strand breaks and that the suppression of this inappropriate repair could rescue the FA cellular phenotypes. Suppression of NHEJ has been shown to relieve sensitivity to ICL-inducing agents in a variety of FA-deficient model systems. In *Caenorhabditis elegans* with a germline defects in *fcd-2* (*FANCD2),* sensitivity to crosslinking agents and abnormal meiotic crossovers were ameliorated after the deletion of lig-4 (Adamo et al., 2010). Depletion of KU80 in human FANCD2 or FANCC deficient patient cells resulted in a partial rescue of ICL hypersensitivity (Adamo et al., 2010). Furthermore, treatment with a DNA-PKcs inhibitor fully or partially alleviated crosslink sensitivity in Hela cells depleted of FANCA or FANCD2 and in *Fanca* or *Fancc* mutant mouse embryonic fibroblasts (MEFs) (Adamo et al., 2010).

In contrast, other studies concluded that the lack of the NHEJ pathway did not rescue the FA defects (Bunting et al., 2012; Houghtaling et al., 2005). These studies showed that knockout of Ku80, DNA-PKcs, or 53bp1 did not rescue ICL sensitivity in MEFs lacking Fancd2 (Bunting et al., 2012; Houghtaling et al., 2005). In a separate paper, a knockout of *Ku70*, but not *Lig4* in FANCC deficient DT40 chicken cells decreased cellular sensitivity and removed chromosomal abnormalities suggesting that it might not be NHEJ per se but the interference of KU heterodimer binding to the inappropriately processed lesions that contributes to the FA cellular phenotype (Pace et al., 2010).

This apparent discrepancy in the outcomes of NHEJ inhibition in the absence of the FA pathway, prompted us to more thoroughly test the consequences of NHEJ inhibition in cell lines derived from Fanconi anemia patients. The use of isogenic controls and direct comparison with complemented cells allows assessment of the extent of any identified rescue and facilitates mechanistic studies. If the inhibition of NHEJ pathway could rescue ICL repair defect in human cells, one would consider NHEJ pathway inhibitors as therapeutic interventions when hematopoietic stem cell transplantation had to be delayed or it was not a viable option in an FA patient. One would then have to understand if such repair in the absence of FA and NHEJ pathways was mutagenic to avoid a possibility of accelerating tumorigenesis during treatment. At a more fundamental level, one would also like to know how the ICL damage is repaired without the FA pathway. Such new pathway, if it existed, would be a target for cancer therapy since its inhibition should synergize with ICL-inducing agents like cisplatin.

To start approaching these questions, we disrupted NHEJ repair pathway with chemical inhibitors, RNA interference (RNAi) and CRISPR-Cas9 in a cell line with most commonly mutated Fanconi gene, *FANCA.* We show that the inhibition of NHEJ pathway by any of the above approaches could not rescue the cellular hypersensitivity of human FA patient cells to ICL damage-inducing agents. Only 53BP1 knockout resulted in a partial rescue of chromosomal instability but no cellular resistance to ICL-inducing agent.

## RESULTS AND DISCUSSION

### Inactivation of DNA-PKcs does not rescue ICL-induced proliferation defects in FA cells

FANCA is the most commonly mutated gene among FA patients, with FANCA mutations accounting for approximately 65% of all FA patient mutations (Kutler et al., 2003). Without FANCA protein, the FA core complex necessary for ubiquitination of FANCD2 and FANCI is not functional and the FA pathway of ICL repair is deficient. Based on these facts, we chose to inactivate NHEJ factors in a human cell line (RA3087) with biallelic deletions of FANCA and not expressing FANCA (Kim et al., 2013). Depletion of DNA-PKcs in RA3087 cells with three independent hairpins (Figure 1A) was sufficient to suppress NHEJ activity as determined by sensitization of RA3087 cells to ionizing radiation (IR) (Figure 1B). Despite NHEJ pathway deficiency, these cells continued to be sensitive to MMC (Figure 1C). Chemical inhibition of DNA-PKcs activity with NU7026 and NU7441 did not rescue MMC sensitivity of RA3087 cells either, despite adequate inhibition of DNA-Pkcs as shown by dosage-dependent IR-induced proliferation defects (Figure 1D-F). DNA-PKcs inhibitor NU7026 treatment also failed to rescue the MMC sensitivity of FA patient cells lacking FANCD2 or SLX4, FA proteins that act downstream of FANCA (Figure 1G).

**Figure 1.**
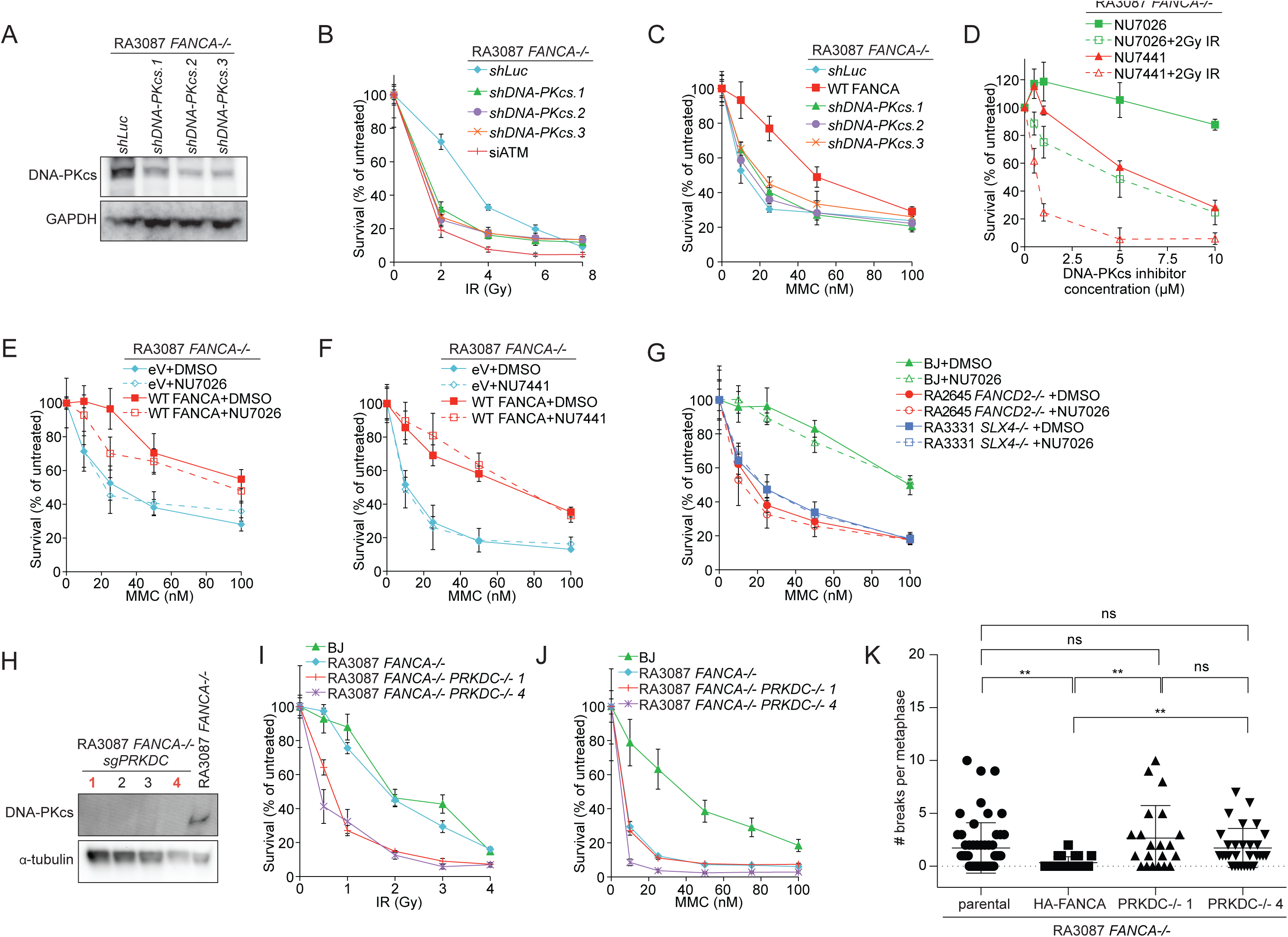
Inactivation of DNA-PKcs fails to restore resistance to MMC in *FANCA*-deficient patient cells. (A) Immunoblot showing the expression of DNA-PKcs in the shRNA transduced *FANCA*-deficient RA3087 patient cells used in (B and C). (B) Survival of RA3087 cells stably transduced with shRNA targeting Luciferase (control) or DNA-PKcs after γ-irradiation (IR) treatment. RA3087 cells with siRNA treatment against ATM was used as IR sensitive-control. Error bars, s.d.. (C) Survival of RA3087 cells stably transduced with shRNA targeting Luciferase (control) or DNA-PKcs and RA3087 cells complemented with FANCA after MMC treatment. Error bars, s.d.. (D) Survival of RA3087 cells after 2 Gy IR and pretreatment with increasing doses of NU7026 or NU7441. Error bars, s.d.. (E) Survival of RA3087 cells stably transduced with empty vector or with FANCA after MMC treatment. Cells were pretreated with 10 μM NU7026 or DMSO. Error bars, s.d.. (F) Survival of RA3087 cells stably transduced with empty vector or with FANCA after MMC treatment. Cells were pretreated with 1 μM NU7441 or DMSO. Error bars, s.d.. (G) Survival of wild type fibroblasts (BJ), RA2645 *FANCD2*-deficient patient cells, RA3331 *SLX4*-deficient patient cells after MMC treatment. The cells were pretreated with 10 μM NU7026 or DMSO. Error bars, s.d.. (H) Immunoblot assessing expression of DNA-PKcs in RA3087 *FANCA*-/- cells and the RA3087 *FANCA-/-DNA-PKcs*-/- clones obtained by CRISPR/Cas9 gene editing. (I) Survival of wild type fibroblasts (BJ), RA3087 *FANCA*-/-, and 2 clones (1 and 4) of RA3087 *FANCA-/-DNA-PKcs*-/- cells after IR. Error bars, s.d.. (J) Survival of wild type fibroblasts (BJ), RA3087 *FANCA*-/-, and 2 clones (1 and 4) of RA3087 *FANCA-/-DNA-PKcs*-/- cells after MMC treatment. Error bars, s.d.. (K) Quantification of chromosomal aberrations after treatment with 50nM MMC for 24 hr. Experiment shown in Figure 1K and 2F were performed in one assay so the same parental and HA-FANCA controls are shown in the graph in both figures.

To eliminate the possibility that the residual DNA-PKcs activity resulted in lack of rescue, we generated knockouts of *PRKDC,* the gene that encodes DNA-PKcs, in RA3087 using CRISPR-mediated genome editing. We derived several *FANCA-/-PRKDC-/*- clones as confirmed by the absence of DNA-PKcs expression at the protein level and by Sanger sequencing at the *PRKDC* genomic locus (Figure 1H, S1A). Two subcloned *FANCA-/-PRKDC*-/- cell lines were confirmed to have mutations predicted to result in the loss of the start codon of DNA-PKcs (Figure S1A). As expected, these *FANCA-/-PRKDC*-/- cell lines were hypersensitive to IR compared to wild type control (BJ) or the parental RA3087 cells (Figure 1I). Nevertheless, both *FANCA-/-PRKDC*-/- clones remained sensitive to MMC (Figure 1J) and continued to show genomic instability after MMC treatment (Figure 1K). Based on these genetic experiments, we conclude that DNA-PKcs inhibition is not sufficient to rescue defective ICL repair in FA patient cells.

### Inactivation of DNA Ligase IV or depletion of Ku70/Ku80 does not rescue ICL damage- induced proliferation defects in FA cells

We next examined the potential role of DNA Ligase IV in suppressing FA phenotypes. LIGIV catalyzes ligation of the processed DNA ends during classical NHEJ (Grawunder et al., 1997). shRNA-targeting of *FANCA* in *LIG4+/+*, *LIG4+/-,* or *LIG4*-/- HCT116 cell lines (Oh et al., 2013) resulted in significant reduction in FANCA protein level and a disappearance of FANCI monoubiquitination in all *shFANCA*-transduced HCT116 cell lines tested (Figure 2B). In this setting, *LIG4* genotype had no effect on the cellular sensitivity to MMC (Figure 2A). We have noted; however, that inactivation of a single (*LIG4*+/-) or both copies (*LIG4*-/-) of *LIG4* without FANCA depletion resulted in dose dependent sensitization of these cells to MMC. It remains to be determined if this is HCT116-specific phenomenon.

**Figure 2.**
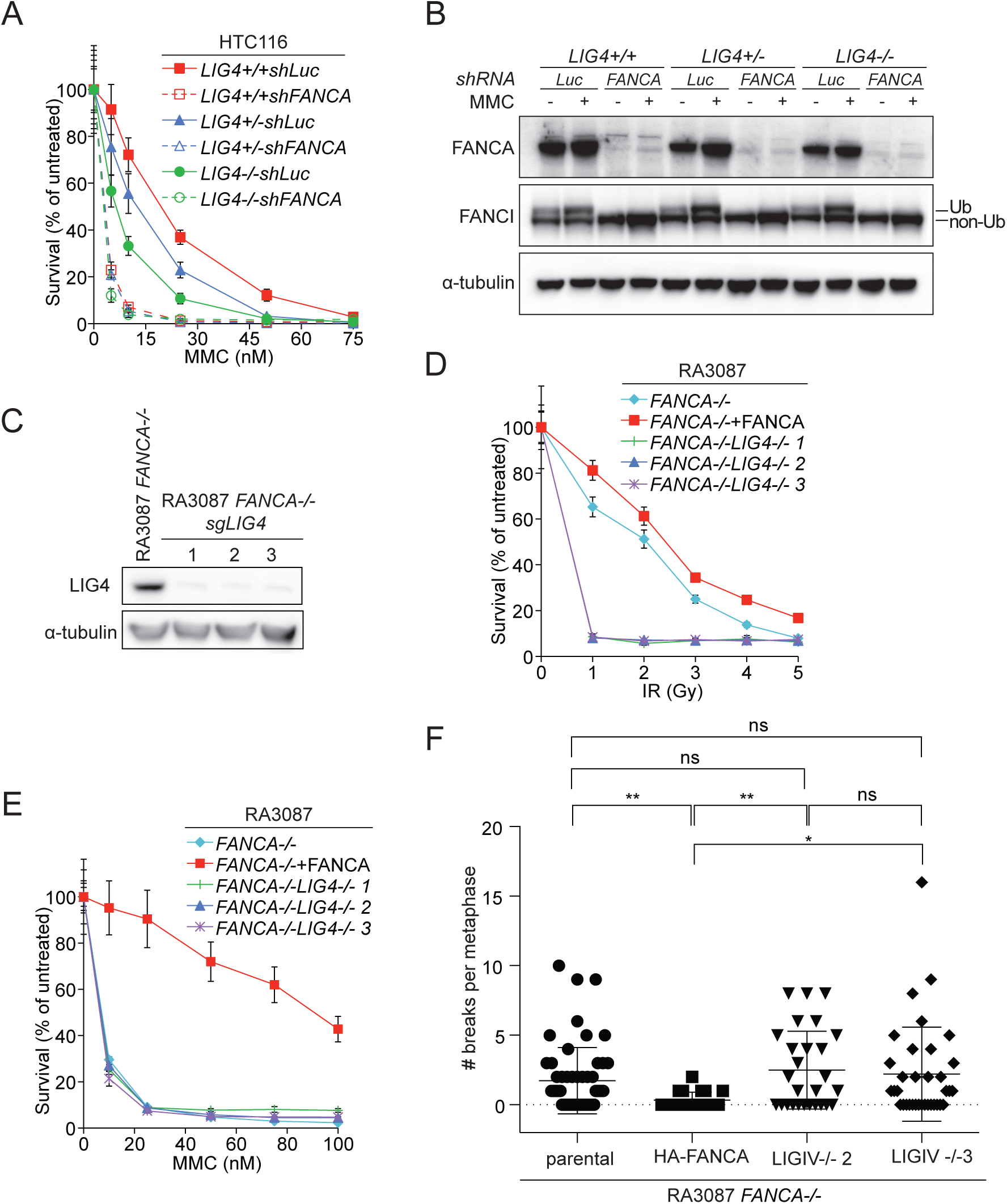
Inactivation of LIG4 does not rescue MMC sensitivity in *FANCA-*deficient patient cells. (A) Survival of *LIG4+/+*, *LIG4*+/-, and *LIG4*-/- HCT116 cells stably transduced with shRNA targeting luciferase sequence (control) or FANCA after MMC treatment. Error bars, s.d.. (B) Immunoblot showing expression of FANCA and FANCI in the shRNA cells used in (A) with or without 24 hr 1 μM MMC treatment. (C) Immunoblot assessing expression of DNA Ligase IV in RA3087 *FANCA*-/- cells and the three clones of RA3087 *FANCA-/-LIG4*-/- cells (1, 2, and 3) obtained by CRISPR/Cas9 gene editing. (D) Survival of RA3087 *FANCA*-/-, RA3087 *FANCA*-/- complemented with FANCA, and 3 clones of RA3087 *FANCA-/-LIG4*-/- cells (1, 2, and 3) after IR treatment. Error bars, s.d.. (E) Survival of RA3087 *FANCA*-/-, RA3087 *FANCA*-/- complemented with FANCA, and 3 clones of RA3087 *FANCA-/-LIG4* -/- cells (1, 2, and 3) after MMC treatment. Error bars, s.d.. (F) Quantification of chromosomal aberrations after treatment with 50nM MMC for 24 hr. Experiment shown in Figure 1K and 2F were performed in one assay so the same parental and HA-FANCA controls are shown in the graph in both figures.

To corroborate the negative results from HCT116 cells, we generated *FANCA-/-LIG4*-/- double knockout RA3087 cells. The knockout of LIG4 was confirmed in three independent *FANCA-/-LIG4*-/- clones at the protein level (Figure 2C) and by their sensitivity to IR (Figure 2D). Even in knockout setting, loss of DNA Ligase IV could not alleviate MMC sensitivity (Figure 2E) or chromosomal abnormalities (Figure 2F).

In the DT40 cells, suppression of *Ku70*, but not *Lig4* suppresses FA phenotypes (Pace et al., 2010). Complete knockout of KU70 (XRCC6) and KU80 (XRCC5) is not attainable in human cells due to the essential function of the KU proteins at telomeres (Fattah et al., 2008; Li et al., 2002), thus we used shRNAs to deplete the KU dimer. KU70 or KU80 depleted cells grew very slowly as expected (Figures S2A-D), but they continued to be sensitive to MMC (Figure S2E).

### Inactivation of DSB repair choice factor, 53BP1, does not rescue ICL-induced proliferation defects but partially suppresses chromosomal abnormalities in FA cells

53BP1 functions to promote NHEJ repair over the competing homologous recombination pathway (Zimmermann and de Lange, 2014). There are conflicting reports on whether 53BP1 inactivation rescues ICL-repair defect in Brca1-deficient mouse cells (Bouwman et al., 2010; Bunting et al., 2012). Notably, 53BP1 deletion did not rescue ICL-induced chromosomal aberrations in Fancd2-deficient MEFs (Bunting et al., 2012). To test the genetic interaction of FANCA and 53BP1, we knocked out 53BP1 in RA3087 cells (Figure 3A and S1). In agreement with published data, we did not observe a significant IR sensitivity in 53BP1-deficient RA3087 cell line (Figure 3C) (Chapman et al., 2012; Iwabuchi et al., 2006; Nakamura et al., 2006; Riballo et al., 2004; Ward et al., 2004). The 53BP1 deletion also did not lead to rescue of cellular MMC sensitivity in human FANCA deficient cells (Figure 3D), although it partially suppressed the chromosomal instability after MMC treatment (Figure 3E). We conclude that the promotion of homologous recombination by deletion of 53BP1, may improve the stability of the genome in FA patient cells but this improvement does not translate into rescue of the cellular sensitivity. This suggests that the instability is not the only determinant of sensitivity to ICL-inducing agents.

**Figure 3.**
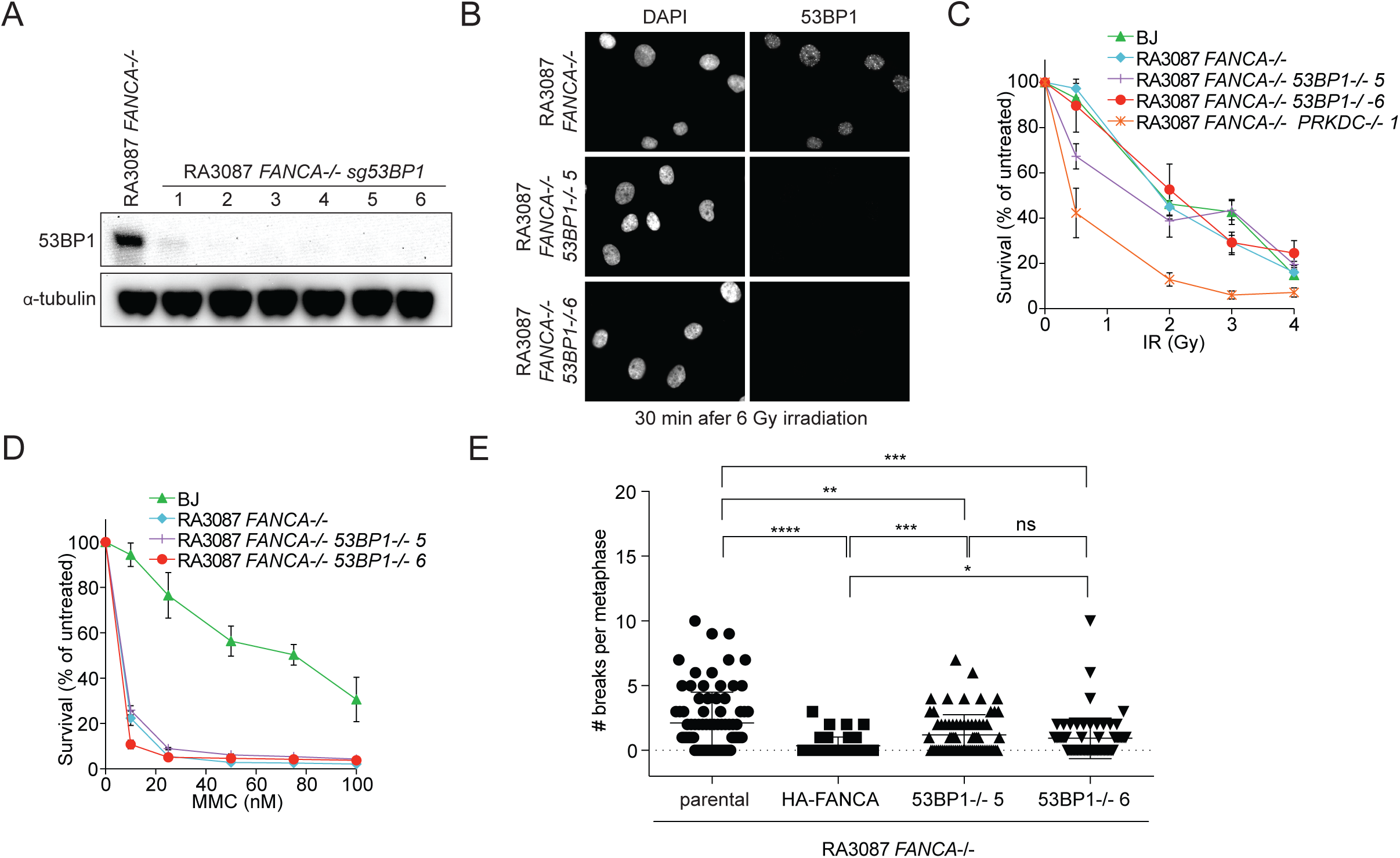
53BP1 inactivation does not rescue MMC sensitivity in *FANCA-*deficient patient cells. (A) Immunoblot assessing expression of 53BP1 in RA3087 *FANCA*-/- cells and the RA3087 *FANCA-/-53BP1*-/- clones. (B) Representative images of 53BP1 foci formation in RA3087 *FANCA*-/- cells and the RA3087 *FANCA-/-53BP1*-/- clones (5 and 6), 30 min following 6 Gy IR. (C) Survival of wild type fibroblasts (BJ), RA3087 *FANCA*-/-, and 2 clones (5 and 6) of RA3087 *FANCA-/-53BP1*-/- cells after IR treatment. RA3087 *FANCA*-/-*DNA-PKcs*-/- 1 CRISPR-targeted clone was used as IR sensitive-control. Error bars, s.d.. (D) Survival of wild type fibroblasts (BJ), RA3087 *FANCA*-/-, and 2 clones (5 and 6) of RA3087 *FANCA-/-53BP1*-/- cells) after MMC treatment. Error bars, s.d.. (E). Quantification of chromosomal aberrations after treatment with 50nM MMC for 24 hr.

Overall, our data show that inhibition of NHEJ is insufficient for providing the rescue of cellular phenotypes associated with Fanconi anemia in human cells. Majority of the experiments in this study were performed using cells from a patient with biallelic mutations in FANCA, the most commonly mutated gene in FA whose presence is necessary for proper function of the whole FA core complex. In addition, FANCD2 and SLX4-deficient cells showed identical results in DNA-PK inhibitor assays. The advantage of our experiments over previous studies performed in human cells is complete knockout of DNAPKcs, Ligase 4 and 53BP1. Use of isogenic cell lines allowed us a direct comparison of cellular behaviors. Our results agree with the genetic results obtained in the mouse models showing lack of rescue of FA phenotypes by inhibition of NHEJ (Bunting et al., 2012; Houghtaling et al., 2005).

The reasons for a discrepancy between conclusions reached in different experimental systems are unclear. However, we note that slow-growing cells often appear resistant to drugs like MMC that act in S phase, so any growth differences between cell lines may lead to differential sensitivity to MMC. Presence of other genetic changes in the cell lines tested or simply differently wired pathways in the case of *C. elegans* or DT40 cells may give different results during interrogation of pathway interactions.

There is a caveat to our study in that the experiments presented here were performed using human patient fibroblasts rather than the hematopoietic stem cells that are directly relevant to FA-associated bone marrow failure. Recently, inhibition of TGF-β signaling was shown to promote resistance to ICL-inducing agents in FA patient cell lines and suppress hematopoietic stem cell (HSC) defects in FA mouse and human systems. This inhibition was correlated with up-regulation of factors involved in homologous recombination and down-regulation of NHEJ factors (Zhang et al., 2016) and the authors proposed that the NHEJ pathway downregulation was responsible for the hematopoietic protection. Given our results in patient fibroblasts and the mouse studies (Bunting et al., 2012; Houghtaling et al., 2005), results of direct inactivation of the NHEJ components should be tested in HSCs of FA patients and mouse models of FA deficiency.

## EXPERIMENTAL PROCEDURES

### Cell culture

Human patient fibroblasts we collected as part of IRB-approved International Fanconi Anemia Registry at the Rockefeller University. Cells were transformed by HPV16 E6E7 and immortalized by expression of catalytic subunit of human telomerase (hTERT). Cells were cultured in 3% oxygen and maintained in DMEM containing 15% FBS, 100 U/mL penicillin, 0.1 μg/mL streptomycin, 0.1 mM nonessential amino acids, and glutamax (Life Technologies).

### Cell survival and cell proliferation analyses

Cell proliferation (KU depletion experiments) was determined using the 96-well plate format. Cell viability was measured daily for 6 days and normalized to the cell viability measured on day 1. Analyses of cell survival under DNA damaging agent treatment were performed in a 6-well or a 96-well plate formats as previously described (Kim et al., 2013). Experiments done in a 6-well format included all experiments shown in Figures 2 and 3 and Figure 1I and J. All other experiments in Figure 1 and S1 were performed using 96-well plate format. Cells were seeded in triplicate at a density of 3.5 × 10^4^ per well in a 6-well plate or 500 cells per well in a 96-well plate (Opaque White MicrotestTM plate (BD, 353296)) before treatment with MMC 24 hours later. For cell survival assay following ionizing radiation, cells were first irradiated in suspension before being plated out. For the 6-well plate format, cell numbers were determined using the Z2 Coulter Counter (Beckman Coulter) after 6–8 days of culture. For the 96-well plate format, cell survival was determined after 5 days in culture using the Cell Titer-Glo reagent (Promega) according to the manufacturer’s instruction. Cell numbers after treatment was normalized to cell numbers in the untreated sample to give the percentage of survival. In the case of NU7026 or NU7441 pretreatment for inhibiting DNA-PKcs, cells were first pre-treated with 10 μM NU7026 or 1 μM NU7441 for 3 hours before being subjected to MMC treatment in a media containing the corresponding DNA-PKcs inhibitor for the duration of the survival assay.

### Chromosome breakage analysis

Cells were exposed to 50 nM MMC for 24 hr before 2 h treatment with 0.167 μg of colcemid per ml of media. Cells were harvested, incubated in 0.075 M KCl for 15 min at 37°C, and fixed in freshly prepared methanol:glacial acidic acid (3:1). Metaphase spreads were prepared by dropping cells onto wet microscope slides. Slides were air-dried at 40°C for 1 hour before staining with 6% Karyomax Giemsa (Life Technologies) in Gurr buffer (Life Technologies) for 3 min. After rinsing with fresh Gurr buffer and distilled water, slides were fully dried at room temperature and scanned using the Metasystems Metafer. T-test was used to determine the statistical significance. The quantification was blinded.

### shRNA, siRNA and CRISPR/Cas9-genome editing

*DNA-PKcs, 53BP1, KU70* and *KU80* mRNA levels were stably knocked down using pAHM (pALPS-Hygro-miR-30). *FANCA* was stably knocked down using pAHM-UltramiR generated by inserting MluI-HpaI fragments of shFANCA pGipz-UltramiR into pAHM. Cells were transduced with shRNA-expressing lentiviruses and selected for 1 week with 200 μg/mL hygromycin. For siRNA transfection, cells were subjected to reverse transfection followed by forward transfection 24 hours later using RNAiMax (Life Technologies). For CRISPR experiments, RA3087 *FANCA−/−* human fibroblasts were transduced with lentivirus expressing pCW-Cas9 vector encoding a doxycycline-inducible Cas9 expression cassette and selected. pCW-Cas9 was a gift from Eric Lander & David Sabatini (Addgene plasmid # 50661) (Wang et al., 2014). Guides were expressed from pLX-sgRNA (gift from Eric Lander & David Sabatini: Addgene plasmid #50662). Following selection in 6 μg/mL blasticidin, cells were treated with 100 ng/mL doxycycline for 48 h before single cells were plated in 96-well plates. Isolated clones were screened and validated for CRISPR-mediated gene inactivation by immunoblotting. Sequences of shRNAs, siRNA and sgRNAs are listed in Supplementary Table 1.

### Antibodies

Immunoblotting and immunofluorescence were performed using the following antibodies: DNA-PKcs (Clone: 25-4, Thermo Scientific, MS-370-P0), DNA Ligase IV (Cell Signaling, D5N5N), 53BP1 (Novus Biologicals, NB100-304), KU70 (Clone N3H10, Thermo Scientific, MS-329-P0), KU80 (a gift from Hiro Funabiki (Postow et al., 2008), FANCA (Bethyl Laboratories, A301-980A-1), FANCI (antibody raised in-house, no. 589), and α tubulin (Sigma-Aldrich, T9026).

## Author Contributions

S.T., B.A.C. and A.S. conceived the ideas and designed experiments for this study. S.T. performed and analyzed RNAi and inhibitor experiments, B.A.C performed and analyzed CRISPR experiments and F.P.L. performed and analyzed the chromosomal breakage experiments. S.T., B.A.C. and A.S. wrote the manuscript.

## ACKNOWLEDGEMENTS

We thank the members of the Smogorzewska laboratory for comments on the manuscript. We thank Scott Lowe for sharing the shRNA cloning protocol and Amaia Lujambio and Elizabeth Garner for the help with the design of shRNA targeting sequences. We also thank Eric Hendrickson for sharing the *LIG4*-deficient HCT116 cell lines. We thank Abraham Brass for sharing pAHM plasmid and Hiro Funabki for the KU80 antibodies. This research was supported by the Doris Duke Charitable Foundation Clinical Scientist Development Award and RO1 HL120922 from NIH. AS is HHMI faculty scholar.

